# The effect of respiratory gases and incubation temperature on early stage embryonic development in sea turtles

**DOI:** 10.1101/2020.05.13.093963

**Authors:** David Terrington Booth, Alexander Archibald-Binge, Colin James Limpus

## Abstract

Sea turtle embryos at high density nesting beaches experience relative high rates of early stage embryo death. One hypothesis to explain this high dead rate is that there is an increased probability that newly constructed nests are located close to maturing clutches whose metabolising embryos cause low oxygen levels, high carbon dioxide levels, and high temperatures. Although these altered environmental conditions are well tolerated by mature embryos, early stage embryos may not be as tolerant leading to an increase in their mortality. To test this hypothesis, we incubated newly laid sea turtle eggs for a week over a range of temperatures in different combinations of oxygen and carbon dioxide concentrations and assessed embryo development and death rates. We found that gas mixtures of decreased oxygen and increased carbon dioxide, similar to those found in natural sea turtle nest containing mature embryos, slowed embryonic development but did not influence embryo mortality of early stage embryos. In contrast, high incubation temperature not only decreased embryo development rate, but prolonged incubation at 34°C was fatal.

## Introduction

The oxygen limitation hypothesis predicts that decreased oxygen availability in the environment and /or limitations in internal oxygen transport can limit aerobic metabolic processes at the intracellular level and thus limit cellular metabolism [1]. In ectothermic animals this potential limitation on cellular process may be exacerbated by high temperatures because high temperatures within the viable temperature range accelerate biochemical reactions and thus oxygen demand, a phenomenon termed the oxygen-capacity-limited thermal tolerance hypothesis [2]. For example, hypoxia increases of lizard embryo mortality at high but not low incubation temperatures [3].

Sea turtle embryos experience varying oxygen availability and incubation temperatures during development in natural nests. Embryos begin their development inside the oviduct of the female where the oxygen partial pressure is very low, less than 1 kPa and embryo development stalls at the mid to late gastrula stage due to lack of oxygen [4]. Once the eggs are laid, they are exposed to higher oxygen partial pressures so that the oxygen limitation is lifted and the embryos break developmental arrest and recommence development [4]. However, once development arrest is broken, if embryos are re-exposed to extremely low oxygen partial pressures they die of asphyxiation [5]. Typically, during the early stages of incubation, sea turtle embryos experience respiratory gases (oxygen and carbon dioxide) partial pressures close to that of the atmosphere above the sand [6,7]. However, during the second half of incubation, as the embryos grow rapidly in size, the combined metabolism of the entire clutch causes the oxygen partial pressure within the nest to decrease and the carbon dioxide partial pressure to increase [6,7]. In green turtle (*Chelonian mydas*) nests, during peak metabolism, the oxygen partial pressure can fall to 10 kPa and carbon dioxide partial pressure climb to 8 kPa [6], while in loggerhead turtle (*Caretta caretta*) nests the equivalent values are 15 - 17 kPa and 3 - 5 kPa, respectively [6,7]. In natural nests the respiratory gases (oxygen and carbon dioxide) always change as mirror images of each other, i.e., as oxygen is consumed by the embryo it released carbon dioxide so that as oxygen decreases within the nest, carbon dioxide increases [6,7,]. The combined metabolism of embryos also generates considerable heat which causes the nest temperature to rise between 2-3°C above the surrounding sand temperature during the latter part of incubation [8,9].

Clutches of eggs laid at high-density sea turtle nesting aggregations such as green turtles on Raine Island, Australia, and olive ridley turtles (*Lepidochelys olivacea*) nesting in arribadas aggregations in Costa Rica, typically experience much higher mortality of embryos than clutches of eggs laid at low-density nesting aggregation beaches [10,11,12,13]. Much of this mortality occurs due to clutch destruction by subsequent nesting females digging up previously constructed nests during their nesting process [13]. However, even in nests that remain undisturbed throughout incubation, embryo mortality is typically much higher than in nests constructed at low-density nesting beaches, and the majority of embryos die at an early stage of development [13]. The physical attributes of the nest environment have been hypothesised as the cause of this elevated embryo mortality [10,11,12,13]. Nests constructed at these high-density beaches may experience lower oxygen, higher carbon dioxide and higher temperature conditions compared to low nest density beaches [10,11,12,13]. Although late stage sea turtle embryos are tolerant to exposure to decreased oxygen, elevated carbon dioxide and elevated temperatures [14,15,16], early stage embryos may not be tolerant, possibly because they have not yet developed the ability to induce a heat-shock protein response which has protective effects against environmental induced stress. Hence, if a newly laid clutch of eggs is laid adjacent to a maturing clutch as may frequently occur at high-density nesting beaches, the newly laid eggs may be exposed to low oxygen, high carbon dioxide and elevated temperature conditions which could increase the frequency of early embryo death. Indeed, high early stage embryo death has been reported in green turtle nests that were laid near maturing nests, however, it was not possible to determine if the respiratory gas conditions or elevated temperatures or a combination of these two factors were responsible for the increase in embryo death [13]. In the current study, through a set of controlled incubation experiments, we investigate if sub-optimal respiratory gases, elevated temperatures or a combination of these factors can cause early embryo death in sea turtle embryos.

## Methods

### Ethics statement

This study was approved by the University of Queensland NEWMA animal ethics committee (certificate SBS/396/18), and eggs were collected under a scientific purposes permit issued by the Queensland Government National Parks Service (permit PTU18-001406).

### Sources of eggs

Loggerhead turtle (*Caretta cartetta*) eggs were collected during oviposition at Mon Repos beach (24.8059°S, 152.4416°E) between 4/12/2019 and 15/12/2019 and green turtle (*Chelonia mydas*) eggs were collected at Heron Island (23.4423°S, 151.9148°E) between 06/01/2019 and 19/01/2019. We performed all experiments at the research facility located within the Mon Repos Conservation Park within 100 m of Mon Repos beach. We transferred eggs from loggerhead turtle nests on Mon Repos beach to the laboratory by hand-carried bucket immediately after collection to be processed. We transferred green turtle eggs immediately after oviposition into an insulated plastic “esky” (60 cm x 35 cm x 35 cm, LxWxH) with its lid open and held overnight in a cool room at 5°C to 8°C. This temperature slows down the embryo developmental rate significantly so that the eggs do not undergo movement-induced mortality during transportation [17,18]. One hour prior to boat departure from Heron Island the esky lid was closed and transported by a 2-hour boat trip to Gladstone followed by a 2.5-hour car trip to Mon Repos where we processed eggs before placing them into incubators, 16 hours after we collected the eggs. We collected two clutches of eggs each night for each species, and we repeated this process once for each species. Hence, in total, we used four clutches of loggerhead turtle eggs and four clutches of green turtle eggs in our experiments.

### Egg incubation

Once at Mon Repos laboratory, we rinsed each egg briefly in distilled water to remove grains of sand, ensuring not to submerge eggs for more than a minute. We weighed eggs and labelled them with a unique identification including clutch and egg number using a 2B pencil. We then placed eggs into a gas tight container labelled with its temperature and respiratory gas mixture treatment. Each container contained ∼8 eggs from one female and ∼8 from the other (no. of eggs in each container depended on total number of eggs in a clutch). Within the containers, we buried eggs in sand sourced from Raine Island that had been heat sterilized. We added distilled water (60g distilled water to 1kg of sand – 6% w/w) to the sand to ensure eggs remained well hydrated during incubation. We placed containers into their designated incubator with unsealed lids exposing eggs to room air for the initial 36 hours of incubation. This procedure insured embryos broke developmental arrest before we exposed eggs to the respiratory gas treatments. After this initial 36 h period, we examined eggs to check if development had begun as indicated by the appearance of a white-patch on top of the egg. In turtle and crocodile eggs, a white-patch occurs in the eggshell immediately above the embryo12-36 hours after the start of incubation as fluid is extracted from the eggshell, causing it to dry and turn white [19,20]. As embryonic development continues, it is thought that dehydration of the albumen below the shell caused by transfer of water from the albumen to the sub-embryonic fluid that surrounds the embryo causes expansion of the white-patch until eventually the white-patch completely surrounds the entire egg [20]. Thus, the rate of expansion of the white-patch is assumed to reflect the rate of embryo development. We assumed that any eggs in which a white-patch was not visible after the initial 36 hours of incubation in air were dead and removed them from the container. We dissected these dead eggs, and examined their contents under a dissecting microscope in order to assign the embryo to a development stage based on morphological criteria [19]. In developing eggs, we traced the outside edge of the white-patch using a 2B pencil and photographed them from above.

After the 36 h initiation period, we returned the eggs to their incubators and applied the gas treatments. For loggerhead turtle eggs, 12 treatments were used, a Latin square design of three incubation temperatures, 27°C, 30°C and 33°C, and four respiratory gas mixtures, O_2_ = 21%, CO_2_ = 0% (room air = control); O_2_ = 17%, CO_2_ = 4%; O_2_ = 14%, CO_2_ = 7% and O_2_ = 10%, CO_2_ = 11%. Unfortunately, by the time green turtle egg trials were run, the supply of the O_2_ = 17%, CO_2_ = 4% gas mixture was exhausted, so this treatment was replaced with an air and 34°C treatment. We choose these temperatures and respiratory gas concentrations as they reflect the range of conditions typically experienced in natural sea turtle nests [6,7,8,9]. Each incubator held a container ventilated at 50ml/min with each of the experimental gas mixtures supplied from premixed gas cylinders. We measured gas concentrations in each container twice a day by aspirating gas leaving the containers into an O_2_/CO_2_ analyser (Quantek 970, USA) to check that eggs were exposed to the appropriate gas concentrations. We also checked temperature inside each incubator twice daily using calibrated mercury in glass thermometers. We incubated all eggs for a further 7 days after initiation of the gas mixture exposure.

After 7 days of gas exposure we removed containers from incubators and examined the eggs. If the initial white-patch had not expanded in size, we assumed the embryo was dead and removed the egg, which we dissected and staged. If the white-patch had expanded we traced the new white-patch boundary with a 2B pencil and photographed the egg from the side. An estimate of white-patch coverage as a percent of the entire egg surface was made using ImageJ (https://imagej.nih.gov/ij/). To confirm embryos within eggs were alive we selected two eggs at random from each treatment (one from each of the two clutches) to be dissected and staged. With the exception of the last two clutches of green turtle eggs, we relocated the surviving eggs into artificial nests on Mon Repos Beach. For the last two green turtle clutches we incubated, after the 7-day gas exposure period, we placed eggs back in their incubators, but incubation continued in room air rather than in their initially assigned gas mixtures. We assessed these eggs for viability after 13 and 46 days of incubation.

### Statistical analysis

We used Pearson correlation to test for a relationship between relative white-patch area and embryo developmental stage. We arcsin transformed relative white-patch area data before ANOVA analysis. Initially we used a mixed model ANOVA (clutch random factor, incubation temperature and respiratory gas treatment fixed factors) for analysis of relative white-patch area for both loggerhead and green turtle eggs. However, clutch was not a significant factor in either species, so we reduced the ANOVA to a two-factor fixed effect model in the reported results. We used Chi-square tests on the two clutches of green turtle eggs that were incubated for 46 days to test for differences in embryo mortality on day 46 across incubation temperature. Statistical significance was assumed if p < 0.05. We performed all statistical procedures using Statistica© version 13 software.

## Results

### Embryo mortality

Only 9 (5 loggerhead turtle eggs, 4 from one clutch, 1 from another clutch; 4 green turtle eggs, 2 from one clutch and 2 from another clutch) out of 827 eggs (409 loggerhead turtle eggs, 418 green turtle eggs) failed to form a white-patch. Dissections indicated that these eggs were fertile but died in very early stages of development soon after oviposition (developmental stages 6-8). For loggerhead eggs that formed a white-patch, with the exception of one embryo incubated at 27°C in 14% O_2_, 7% CO_2_, there was no embryo mortality between the 36 h to 7 day period of gas exposure across all treatments (Table 1). Similarly, mortality of green turtle embryos was very low across all treatments between 36 h and 7 days of incubation (Table 2).

**Table 1.** Experimental treatments used to incubate loggerhead turtle eggs indicating survival after 7 days of exposure to the incubation treatment.

| Incubation<br>temperature<br>(°C) | Gas treatment | Number<br>of eggs<br>set | Number of<br>eggs with<br>white-patches<br>at 36 h | Number of<br>embryos<br>alive at 7<br>days | Proportion<br>survived (%) |
| --- | --- | --- | --- | --- | --- |
| 27 | 21% O<sub>2</sub>, 0% CO<sub>2</sub> | 32 | 31 | 31 | 100 |
| 27 | 17% O<sub>2</sub>, 4% CO<sub>2</sub> | 34 | 33 | 33 | 100 |
| 27 | 14% O<sub>2</sub>, 7% CO<sub>2</sub> | 36 | 36 | 35 | 97 |
| 27 | 10% O<sub>2</sub>, 11% CO<sub>2</sub> | 32 | 32 | 32 | 100 |
| 30 | 21% O<sub>2</sub>, 0% CO<sub>2</sub> | 32 | 32 | 32 | 100 |
| 30 | 17% O<sub>2</sub>, 4% CO<sub>2</sub> | 32 | 32 | 32 | 100 |
| 30 | 14% O<sub>2</sub>, 7% CO<sub>2</sub> | 32 | 31 | 31 | 100 |
| 30 | 10% O<sub>2</sub>, 11% CO<sub>2</sub> | 32 | 32 | 32 | 100 |
| 33 | 21% O<sub>2</sub>, 0% CO<sub>2</sub> | 36 | 35 | 35 | 100 |
| 33 | 17% O<sub>2</sub>, 4% CO<sub>2</sub> | 35 | 35 | 35 | 100 |
| 33 | 14% O<sub>2</sub>, 7% CO<sub>2</sub> | 40 | 39 | 39 | 100 |
| 33 | 10% O<sub>2</sub>, 11% CO<sub>2</sub> | 36 | 36 | 36 | 100 |

**Table 2.** Experimental treatments used to incubate green turtle eggs indicating survival after 7 days of exposure to the incubation treatment.

| Incubation temperature (°C) | Gas treatment | Number of eggs set | Number of eggs with white-patches at 36 h | Number of embryos alive at 7 days | Proportion survived (%) |
| --- | --- | --- | --- | --- | --- |
| 27 | 21% O<sub>2</sub>, 0% CO<sub>2</sub> | 36 | 36 | 36 | 100 |
| 27 | 17% O<sub>2</sub>, 4% CO<sub>2</sub> | 35 | 35 | 35 | 100 |
| 27 | 10% O<sub>2</sub>, 11% CO<sub>2</sub> | 37 | 37 | 37 | 100 |
| 30 | 21% O<sub>2</sub>, 0% CO<sub>2</sub> | 36 | 36 | 36 | 100 |
| 30 | 17% O<sub>2</sub>, 4% CO<sub>2</sub> | 35 | 35 | 35 | 100 |
| 30 | 10% O<sub>2</sub>, 11% CO<sub>2</sub> | 37 | 37 | 36 | 97 |
| 33 | 21% O<sub>2</sub>, 0% CO<sub>2</sub> | 36 | 35 | 35 | 100 |
| 33 | 17% O<sub>2</sub>, 4% CO<sub>2</sub> | 36 | 36 | 36 | 100 |
| 33 | 10% O<sub>2</sub>, 11% CO<sub>2</sub> | 36 | 35 | 33 | 94 |
| 34 | 21% O<sub>2</sub>, 0% CO<sub>2</sub> | 94 | 92 | 89 | 97 |

Although we observed little embryo mortality during the first 8.5 days of incubation (1.5 days in room air + 7 days in gas mixture treatment), we observed an increase in mortality associated with higher incubation temperatures in the two clutches of green turtle eggs we monitored for a longer period (Fig. 1). No eggs survived incubation at 34°C for a prolonged period, and mortality was significantly increased at 33°C (Fig. 1). Dissections of dead eggs at 46-days indicated blood had been present but embryos had deteriorated so it was not possible to identify their developmental stage at death, but it must have occurred early in development because no signs of a carapace were present.

**Fig 1.**
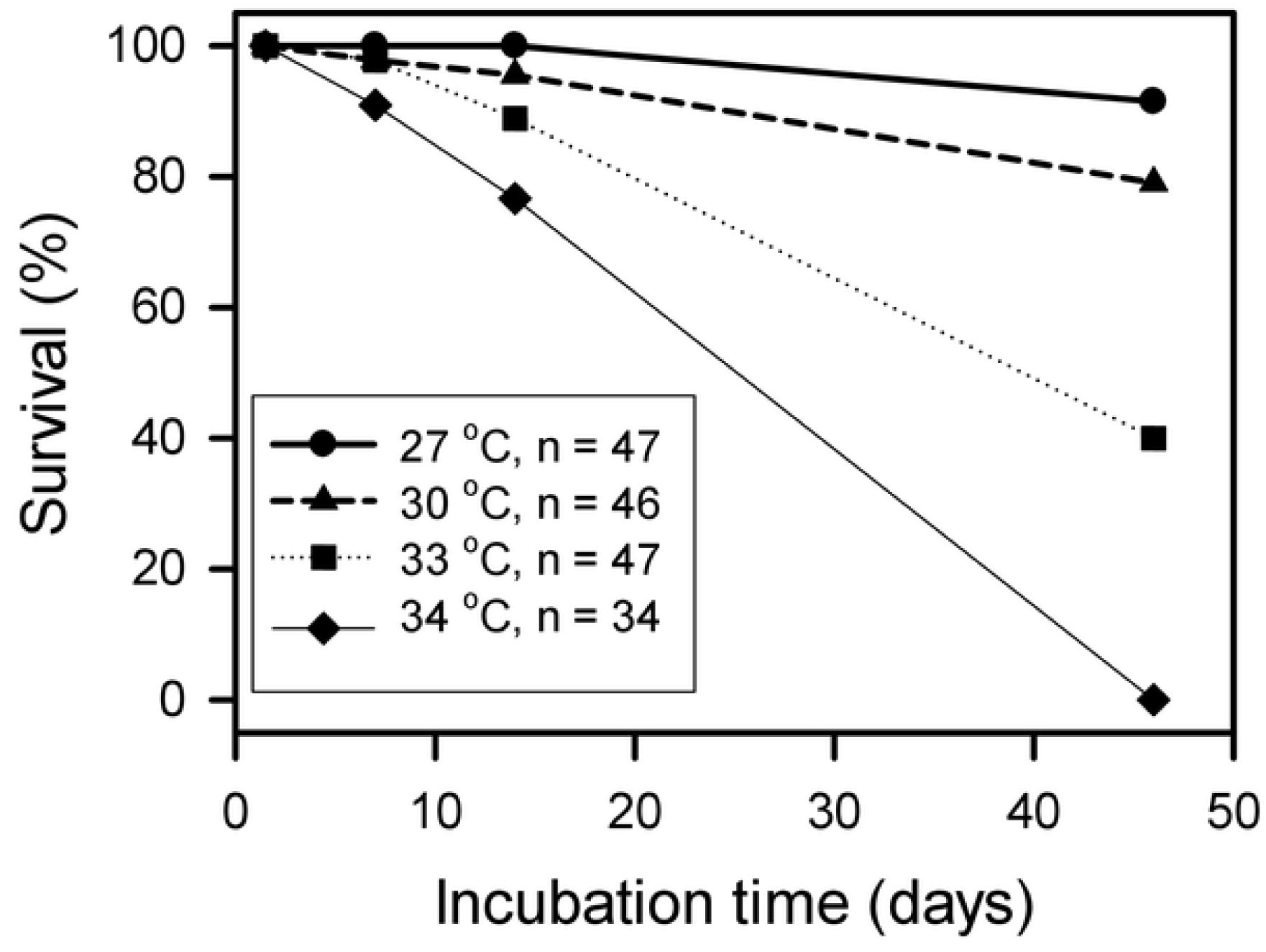
Relationship between incubation temperature and embryo survival during incubation for two clutches of green turtle eggs. For each incubation temperature, data were pooled across gas treatments. Survival on day 46 was significantly reduced at 33°C (□^2^ = 10.16, df = 1, P = 0.001) and 34°C (□^2^ = 65.47, df = 1, P < 0.001) compared to 27°C and 30°C.

### Embryo development

We assessed embryo development after the 7-day exposure to the respiratory gas treatments by two methods; direct staging of embryonic development by dissection of four eggs from each treatment (one egg from each clutch), and by the relative white-patch area. Relative white-patch area was correlated with developmental stage in both loggerhead (r^2^ = 0.33, t = 2.21, p = 0.05, n = 12) and green (r^2^ = 0.62, t = 3.34, p = 0.012, n = 9) turtles indicating that relative white-patch area is a good indicator of developmental stage during early incubation. For this reason, and because the sample size for relative white-patch area was much greater than for developmental stage, only detailed analysis of relative white-patch area is presented. In loggerhead and green turtle embryos, both incubation temperature and respiratory gas treatment influenced growth of the white-patch (Tables 3, 4). The trend in both species was for growth rate to increase from 27°C to 30°C, and then to remain similar between 30°C and 33°C, and for green turtle eggs that were incubated at 34°C, growth of the white-patch was slowest at 34°C (Fig. 2). The rate of growth of the white-patch decreased as the respiratory gases embryos were exposed to deviated further from atmospheric air, for both loggerhead and green turtle embryos (Fig. 3). While dissecting eggs to record embryo development stage, we found that one green turtle embryo from the 33°C, 21% O_2_, 0% CO_2_ treatment was malformed.

**Table 3.** Fixed factor ANOVA table for loggerhead egg arcsin transformed relative white-patch area after 7 days exposure to experimental incubation regimes.

| Effect | SS | DF | MS | F | P |
| --- | --- | --- | --- | --- | --- |
| Intercept | 210 | 1 | 210 | 9383 | < 0.001 |
| Incubation temperature | 1.23 | 2 | 0.62 | 27.4 | < 0.001 |
| Respiratory gas | 15.3 | 3 | 5.13 | 229 | < 0.001 |
| Incubation temperature*respiratory gas | 2.20 | 6 | 0.37 | 16.3 | < 0.001 |
| Error | 8.70 | 388 | 0.02 |  |  |

**Table 4.**
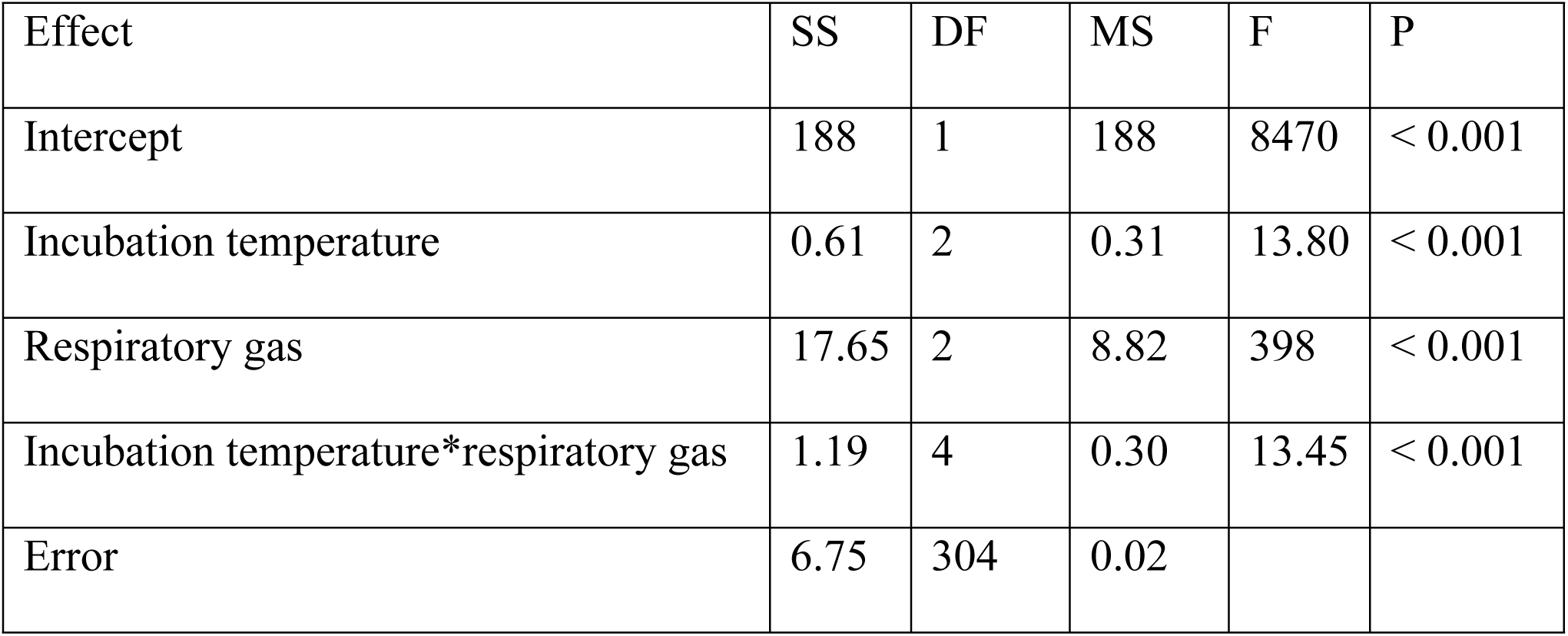
Fixed factor ANOVA table for green egg arcsin transformed relative white-patch area after 7 days exposure to experimental incubation regimes.

| Effect | SS | DF | MS | F | P |
| --- | --- | --- | --- | --- | --- |
| Intercept | 188 | 1 | 188 | 8470 | < 0.001 |
| Incubation temperature | 0.61 | 2 | 0.31 | 13.80 | < 0.001 |
| Respiratory gas | 17.65 | 2 | 8.82 | 398 | < 0.001 |
| Incubation temperature*respiratory gas | 1.19 | 4 | 0.30 | 13.45 | < 0.001 |
| Error | 6.75 | 304 | 0.02 |  |  |

**Fig 2.**
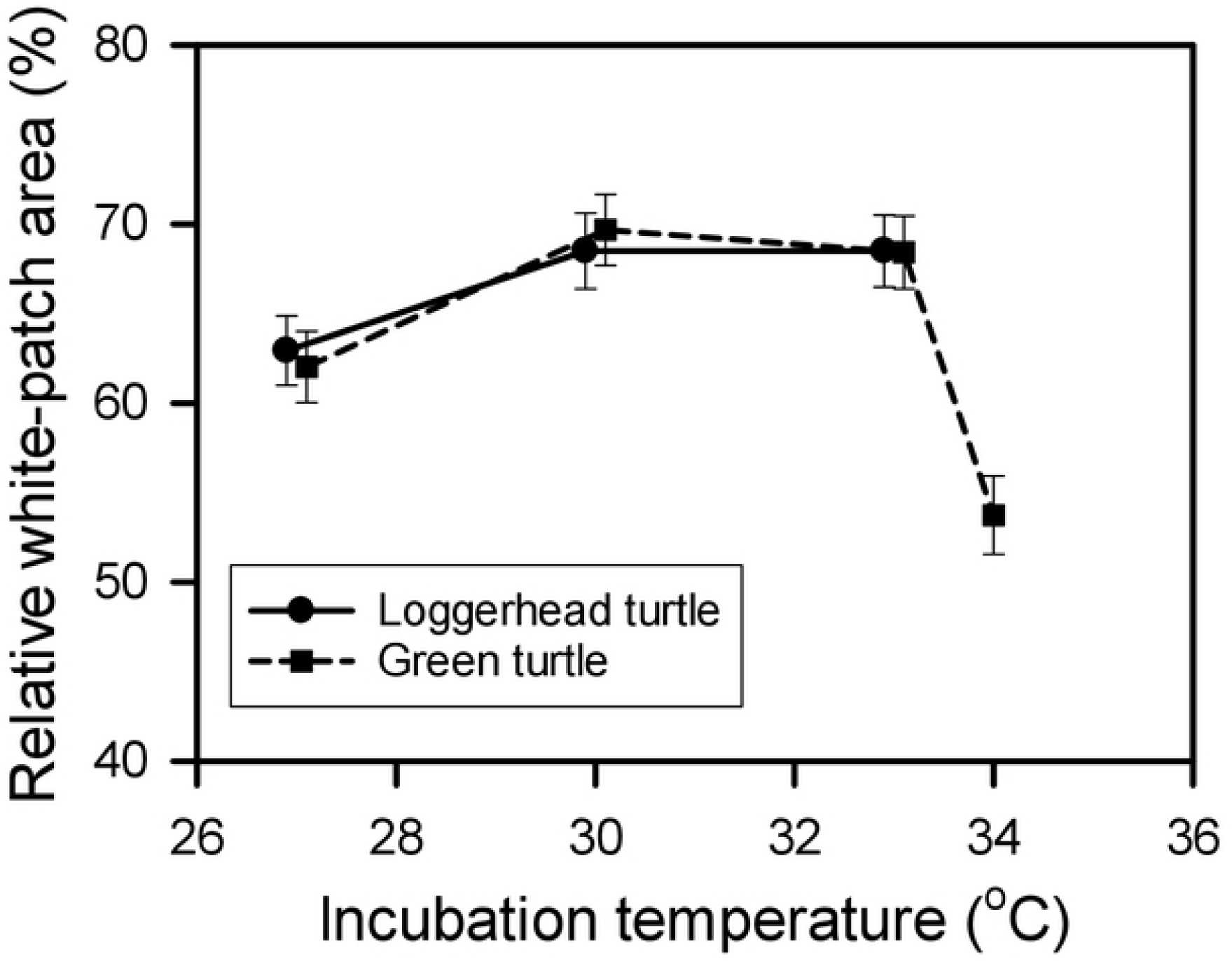
The influence of incubation temperature (pooled across respiratory gas treatments) on relative white-patch area after 8.5 days of incubation for green and loggerhead turtle eggs.

**Fig 3.**
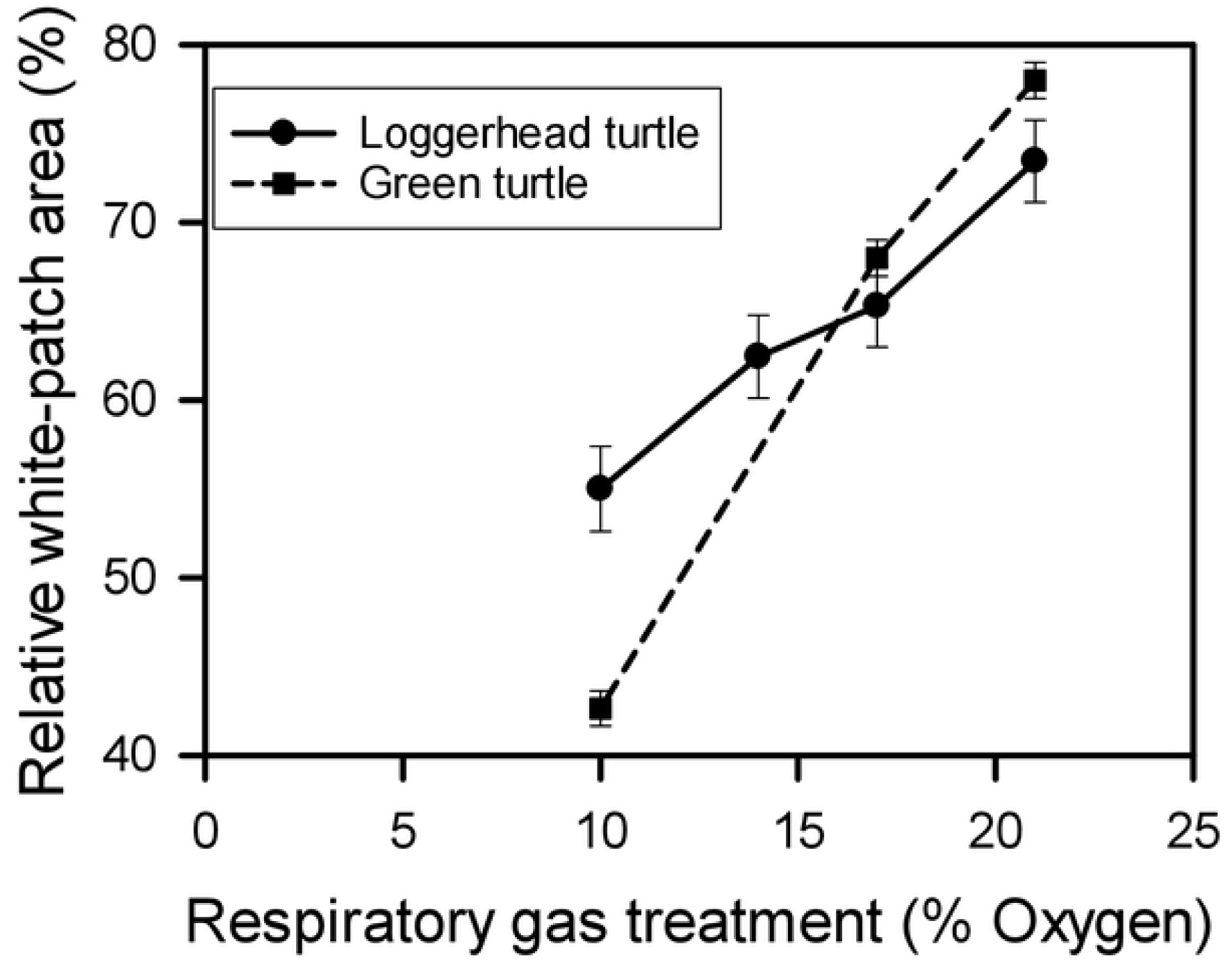
The influence of incubation respiratory gas treatment (pooled across incubation temperatures) on relative white-patch area after 7 days exposure to the respiratory gas treatment for green and loggerhead turtle eggs. Respiratory gas treatments were: 21% O_2_, 0% CO_2_; 17% O_2_, 4% CO_2_; 14% O_2_, 7% CO_2_ (loggerhead turtles only); 10% O_2_, 11% CO_2_.

## Discussion

### Respiratory gases

Contrary to our expectation, exposure of early stage embryos to low oxygen and high carbon dioxide gas mixtures did not influence mortality during the first week of incubation. Hence, we reject the respiratory gas hypothesis as an explanation of early embryo death syndrome so prevalent in high nest density sea turtle rookeries. However, we found that as the respiratory gases deviated further from atmospheric air, the rate of embryo development decreased, so clearly there was a deleterious effect of prolonged exposure to sub-atmospheric respiratory gases at this early stage of embryonic development. If new clutches are laid in close proximity to maturing clutches, this can potentially cause the respiratory gases in the newly laid clutch to deviate significantly from atmospheric air. Under these conditions, the clutch may experience a longer incubation period than other clutches that are not influenced by altered respiratory gases. However, in nature, clutches are only exposed to low oxygen and high carbon dioxide partial pressures during the last 2 weeks of incubation. After this time, hatchlings escape the nest and oxygen and carbon dioxide partial pressures return to air conditions. So, even if a new clutch was laid next to a late maturing clutch, embryos in the new clutch should only be exposed to adverse respiratory gases for a maximum of two weeks.

Our data suggest that early embryo growth rate of green turtles is more sensitive to low oxygen and high carbon dioxide conditions than logger turtle embryos because the relative white-patch area after 7 days of exposure to the most extreme respiratory gas condition was considerable smaller in green turtle eggs. This was an unexpected finding because green turtle clutches are typically buried in deeper nests than loggerhead turtle nests, and as a consequence the in-nest partial pressure of oxygen are typically lower and the partial pressure of carbon dioxide higher in green turtle nests than in loggerhead turtle nests [6,7]. Hence, one might expect green turtle embryos to be less sensitive to altered respiratory gases than loggerhead turtle embryos. It is possible, although unlikely, that the chilling we applied to newly collected green turtle eggs for protection of embryos during transport had a downstream effect on embryo growth rate. If this is the case, it could explain the difference in growth rate between green and loggerhead turtle embryos we observed. However, oxygen limitation is the probable mechanism behind the slowing of embryonic development we observed in both green turtle and loggerhead embryos when exposed to low partial pressures of oxygen. Oxygen limitation would occur if the aerobic metabolic pathways required for embryo growth and development are slowed by a decreased oxygen partial pressure. The slowing of embryonic development when exposed to decreased oxygen partial pressure is a commonly reported phenomenon, having been documented in molluscs [22], fishes [23], amphibians [24], mammals [25], freshwater turtles [26], and sea turtles [27].

### Incubation temperature

Although incubation temperature appeared to have little effect on embryo mortality over the first 8.5 days of incubation, prolonged incubation at 34°C resulted in death of all embryos as has been documented previously for sea turtle embryos [14,28,29,30]. The observation that an embryo was still alive, but malformed after 8.5 days of incubation at 33°C suggests that it is possible that even though embryos incubated at this temperature are still alive after a week of incubation, some embryos have developed high temperature induced teratogenic malformations that will result in embryo death during later incubation. Hence, high nest temperatures experienced early in incubation is a likely explanation for the large number of nests experiencing a high rate of early embryo death syndrome in high-density nesting aggregations. Indeed, a natural green turtle clutch that experienced 80% early embryo death at Raine Island experienced an average temperature of 35.5°C during its first week of incubation [13]. In natural sea turtle nests, a rise in nest temperature due to metabolic heating is always accompanied by a fall in oxygen and increase in carbon dioxide, and it is generally theorised that a decrease in oxygen would exacerbate the detrimental effect of high temperature [2], and this has been indicated in developing lizard embryos [3]. However, we found no evidence of this exacerbated effect in our sea turtle embryos, with high survivability during the first week of incubation in hypoxia across all experimental temperatures.

Although high incubation temperatures were not immediately fatal during the first week of incubation, they did retard the growth of early stage embryos. In ectotherms, including sea turtles, embryonic growth and development rate increase with an increase in incubation temperature within the viable temperature range. This explains the increase in embryo development rate we observed between 27°C and 30°C, however development rate did not continue to increase at 33°C, and actually decreased at 34°C. It would appear that incubation at temperatures of 33°C and higher are sub-optimal and that although development can continue, these high temperatures probably cause cellular damage that needs to be repaired, and this damage slows the rate of development. Ultimately, at temperatures of 34°C and higher, the high temperature induced damage results in embryo malformation and death.

In summary, although we found early stage sea turtle embryos development rate is retarded when exposed to partial pressures of oxygen and carbon dioxide typically encountered in maturing sea turtle cluthes, such exposure is not fatal. In contrast, when we exposed early stage sea turtle embryos to incubation temperatures of 33°C and above, development was slowed, and when exposed to 34°C the embryos died. At nesting beaches that experience high-density nesting, the close proximity of nests means that many newly laid clutches could experience high temperatures due to the metabolic heat production of nearby maturing clutches, and this could result in elevated rates of early embryo death syndrome. Hence, as the number of nesting females increases at high-density rookeries, the number of newly laid clutches exposed to high temperatures increases, and therefore the proportion of clutches experiencing early embryo death syndrome increases as reported for green turtle clutches at Raine Island [13]. A management strategy that could mitigate this increase in early embryo death syndrome could be to increase the beach area suitable for nesting so that nests could be spaced further apart, or other strategies such as shading or irrigation that reduce sand temperature.

## Acknowledgements

This project was funded by the The Raine Island Recovery Project which is a five-year, $7.95 million collaboration between BHP, the Queensland Government, the Great Barrier Reef Marine Park Authority, Wuthathi and Kemerkemer Meriam Nation (Ugar, Mer, Erub) Traditional Owners and the Great Barrier Reef Foundation.

